# Intracellular XBP1-IL-24 axis dismantles cytotoxic unfolded protein response in the liver

**DOI:** 10.1101/658666

**Authors:** Jianye Wang, Bian Hu, Zhicong Zhao, Haiyan Zhang, He Zhang, Zhenjun Zhao, Xiong Ma, Bin Shen, Beicheng Sun, Xingxu Huang, Jiajie Hou, Qiang Xia

## Abstract

Endoplasmic reticulum (ER) stress-associated cell death is prevalent in various liver diseases. However, the determinant mechanism how hepatocytes survive unresolved stress was still unclear. Interleukin-24 (IL-24) was previously found to promote ER stress-mediated cell death, and yet its expression and function in the liver remained elusive. Here we identified an anti-apoptotic role of IL-24, which transiently accumulated within ER-stressed hepatocytes in a X-box binding protein 1 (XBP1)-dependent manner. Disruption of IL-24 increased cell death in the CCL_4_- or APAP-challenged mouse liver or Tm-treated hepatocytes. In contrast, pharmaceutical blockade of eukaryotic initiation factor 2α (eIF2α) or genetical ablation of C/EBP homologous protein (CHOP) restored hepatocyte function in the absence of IL-24. In a clinical setting, patients with acute liver failure manifested a profound decrease of hepatic IL-24 expression, which was associated with disease progression. In conclusion, intrinsic hepatocyte IL-24 maintains ER homeostasis by restricting the eIF2α-CHOP pathway-mediated stress signal, which might be exploited as a bio-index for prognosis or therapeutic intervention in patients with liver injury.

## Introduction

The liver, one of the most vital organs in metabolic homeostasis, has a unique potential to fully recover from acute liver injury. Despite recent studies elucidating various molecular pathways involved in liver damage [1], a further understanding of the pivotal life-and-death decision mechanism is needed to improve current therapeutics. Endoplasmic reticulum (ER) content is rich in hepatocytes and participates in the processes of synthesizing, folding and trafficking of proteins [2, 3]. Environmental stimuli or nutrient fluctuations disrupt the ER protein-folding procedure, referred to as ER stress [4]. With an accumulation of misfolded proteins in ER lumen, the unfolded protein response (UPR), a collection of intracellular signal pathways, is activated to increase protein-folding capacity and reduce global protein synthesis. Once the molecular adaption fails in resolving the protein-folding defect, hepatocytes enter persistent ER stress, which results in apoptosis[5]. ER stress-related apoptosis has been found in fatty liver disease, viral hepatitis, and alcohol or drug induced liver injury [3, 6, 7]. The transcription factor C/EBP homologous protein (CHOP) mediates the most well-characterized pro-apoptotic pathway resulted from unresolved ER stress. CHOP induces the expression of pro-apoptotic BH3-only protein Bim, the cell surface death receptor TRAIL receptor 2, and inhibits Bcl2 transcription [8-11]. As previously reported, CHOP-deficient mice were protected from acetaminophen (APAP)-induced liver damage and conferred a survival advantage [12].

Interleukin-24 (IL-24) was first identified as a negative regulator in human melanocytes[13, 14]. As an IL-10 superfamily member, IL-24 has been reported to exert a bystander anti-cancer function, but has no deleterious effect toward non-cancerous cell [13, 15-17]. Like other secretory proteins, IL-24 precursor, which is 206 amino acids in length, translocates to the ER lumen before it proceeds to the secretory pathway. Independently of its cognate receptors, adenovirus-mediated IL-24 overexpression in melanoma cells led to induction of apoptosis by interaction with glucose-regulated protein 78 (GRP78) and upregulation of GADD family genes, including CHOP [18, 19]. Nonetheless, little is known about IL-24 expression and its correlation with ER stress in non-cancerous cells. Interestingly, IL-24 production was elevated in diabetic pancreatic islets, where it induced beta cell ER stress and impaired glucose tolerance [20]. But it remains unclear whether IL-24 adapts ER homeostasis in epithelial cells. Given the abundant IL-24 expression in the normal mouse or human liver detected in our preliminary experiments, the role of hepatocyte IL-24 in liver diseases has yet to be deciphered. To search for a possible link between IL-24 and ER stress within hepatocytes, we employed two mouse models characterizing IL-24 in the duration of acute liver injury. Unexpectedly, IL-24 deficiency did not alleviate liver damage but sensitized ER stressed hepatocytes to death. By monitoring tunicamycin (Tm)-stimulated mouse hepatocytes *in vitro* or manipulating the IL-24 level or UPR pathway *in vivo*, we further confirmed anti-apoptotic function of intracellular IL-24. Indeed, we revealed that hepatocyte IL-24 governs the intrinsic adaption to ER stress by control of PERK-eIF2α-CHOP pathway. Collectively, these results highlight profound implications for understanding hepatocyte ER homeostasis and identify IL-24 as a critical anti-stress factor in the liver.

## Results

### Hepatocyte IL-24 transiently increases during ER stress-related acute liver injury

Firstly, we detected the expression of IL-24 among different organs in normal wild type (WT) mice and found it was most highly expressed in the liver (Expanded View Fig. 1A). To explore whether liver IL-24 is linked to ER stress, we treated WT mice with a single dose of CCL_4_ (2ml/kg) as reported previously^19^. Serum levels of alanine aminotransferase (ALT) and aspartate aminotransferase (AST) were markedly elevated and peaked at 48 h post treatment, then returned to baseline at 72 h time point (Expanded View Fig. 1B). In context, exposure to CCL_4_ unaffected the ER chaperon GRP78 but tremendously evoked CHOP expression in the liver (Fig. 1A&B). Noticeably, IL-24 mRNA level was transiently increased at 24 h, then decreased and reverted to normal level 72 h post CCL_4_ administration. A same trend was observed in IL-24 protein level (Fig. 1A). Regarding the potential inflammatory responses caused by CCL_4_, we also measured the serum IL-24 protein level. Nonetheless, CCL_4_-treated mice exhibited undetectable serum IL-24, which was comparable to that in none-treated WT or IL-24 KO mice (data not show). Given the fact that hepatocytes take up the majority of hepatic cells, we asked whether the fluctuation of IL-24 expression was occurred in ER stressed-hepatocytes. To answer this question, we investigated IL-24 expression in murine hepatocyte cell line AML12 in the presence of an ER stress inducer Tm. Consistent with the *in vivo* observation, a transient increase of IL-24 level was recapitulated in AML12 upon ER stress, as accompanied by accumulating CHOP expression. These results suggested a potential role of non-secreted IL-24 and prompted us to understand how IL-24 was involved in hepatocyte ER stress.

**Fig. 1:**
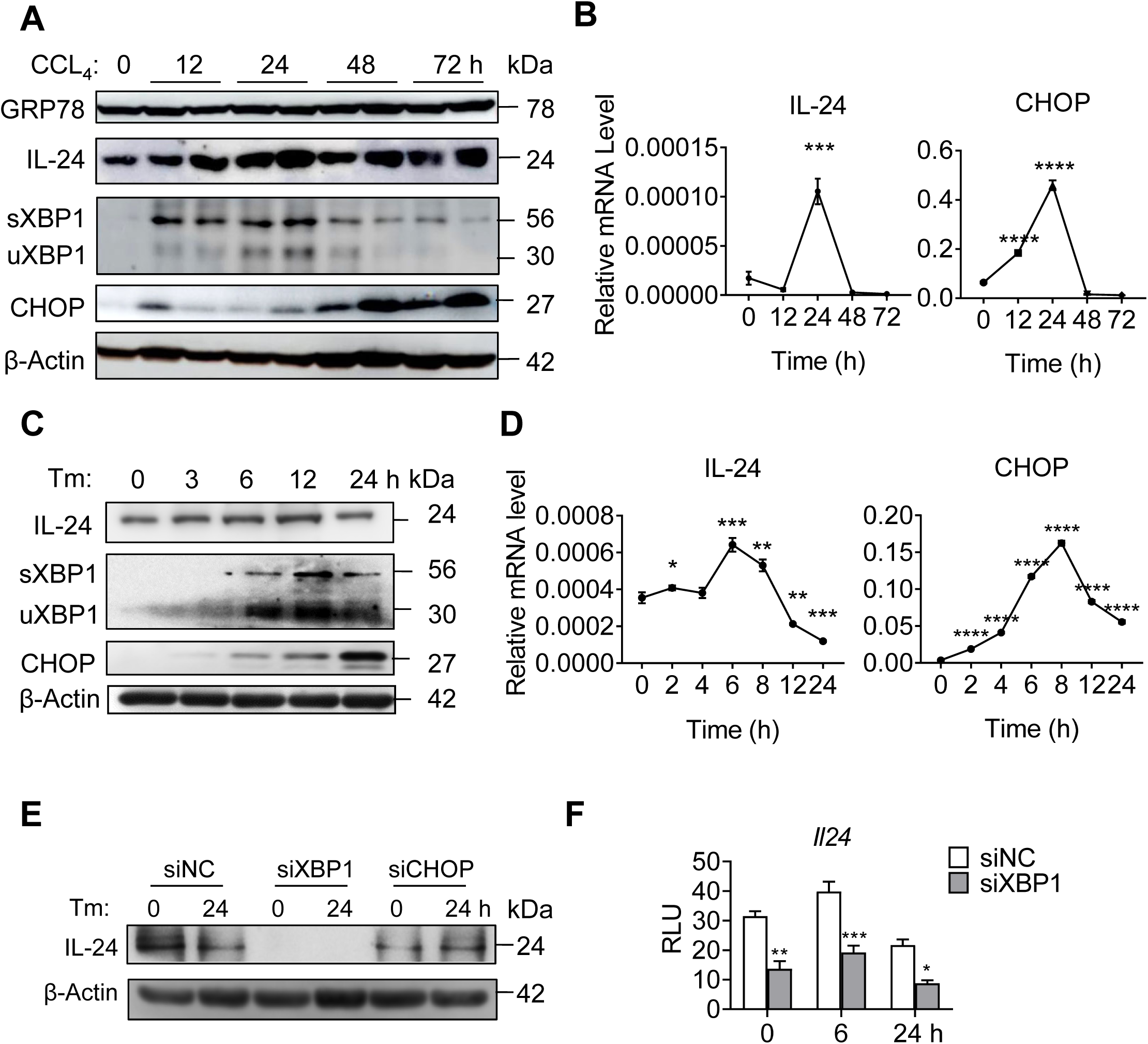
IL-24 expression in the ER stressed-mouse liver and hepatocytes. (A, B) WT mice were intraperitoneally injected with a single dose of CCL_4_ (2ml/kg). (A) GRP78, IL-24 and CHOP protein levels in the liver tissues were evaluated by western blot at indicated time points. Results are normalized to β-actin. *n* = 3 independent experiments. (B) IL-24 and CHOP mRNA levels in the liver tissues were assessed by qRT-PCR at indicated time points. Results are normalized to β-actin. *n* = 3 biological replicates. (C, D) AML12 cells were exposed to Tm (5 µg/ml) for the indicated time period. IL-24 and CHOP levels were evaluated by western blot (C) and qRT-PCR (D) at indicated time points. *n* = 2 independent experiments (C) or 3 biological replicates (D). (E) AML12 cells were transfected with indicated siRNAs or negative control (NC) 24 h prior to Tm treatment. IL-24 protein levels at indicated time points were assessed by western blot. *n* = 3 independent experiments. (F) *Il24* promoter activity in AML12 cells expressing indicated siRNAs, as quantified using luciferase assay. *Renilla* luciferase activity was normalized to firefly activity and presented as relative luciferase activity. *n* = 3 biological replicates. Data are presented as means ± SEM. \**P* < 0.05, \*\**P* < 0.01, \*\*\**P* < 0.001, \*\*\*\**P* < 0.0001. *P*-values were determined by two tailed *t*-test.

The transcription factors activating transcription factor 4 (ATF4), ATF6, sliced X-box binding protein 1 (sXBP1) and CHOP regulate UPR-related gene expression[5]. Like CHOP, other three molecules were also upregulated in the CCL_4_-exposed mouse liver or ER stressed-AML12 cells, among which sXBP1 was the first to peak (Expanded View Fig. 1C&D). Intriguingly, the murine *Il24* promoter harbors conserved binding motifs for ATF6/XBP1 and CHOP [21, 22](Expanded View Fig. 2A). To explore how IL-24 expression was affected in response to ER stress, we transfected AML12 cells with small interfering RNAs (siRNAs) targeting ATF4, ATF6, XBP1 and CHOP prior to Tm stimulation. IL-24 mRNA levels in these siRNA-expressing cells all decreased as compare to the negative control (NC) (Expanded View Fig. 2B), suggesting a regulatory relation between hepatocyte IL-24 and UPR pathways. Importantly, siXBP1 most significantly inhibited IL-24 mRNA level and blocked its upregulation in response to ER stress. Silencing of XBP1 (but not CHOP or ATF6) repressed IL-24 protein level as well as *Il24 promoter* activity in Tm-stimulated AML12 cells (Fig. 1E&F and Expanded View Fig. 2C). Furthermore, we isolated primary hepatocytes from conditional XBP1 KO (Xbp1^f/w^;Alb^Cre^) mice. XBP1 depletion unaffected cell viability under ER stress, but reduced IL-24 in both mRNA and protein levels (Expanded View Fig. 2D-F).

### Hepatocyte IL-24 deficiency promotes ER stress-related liver injury

To further dissect the underlying impact of IL-24 fluctuation, IL-24 KO mice, in which 3306 bp of IL-24 allele was depleted, were subjected to CCL_4_-induced liver injury. In comparison to the WT mice, IL-24-null littermates were more susceptible to CCL_4_-induced liver injury, exhibiting a relatively higher ALT and AST level and a lower survival rate (Fig. 2A&B). The exacerbated liver damage in IL-24-null mice was visualized by hematoxylin and eosin (H&E). Meanwhile, a marked increase in the percentage of terminal deoxynucleotidyl transferase dUTP nick end labeling (TUNEL)-positive hepatocyte was observed in IL-24-null mice with respect to WT mice (Fig. 2C&D). Besides, proliferating cell nuclear antigen (PCNA) staining showed an increase of cell proliferation in IL-24 KO mice (Expanded View Fig. 3A), possibly due to compensatory liver regeneration. We then checked the inflammatory status and detected higher IL1A and IL6 and lower TNFA mRNA expression in the IL-24-deficient mouse liver (Expanded View Fig. 3B). Overdose of APAP, an analgesic and antipyretic drug, is the leading cause of drug-induced acute liver injury [23]. Accordingly, we subjected IL-24-null mice to oral administration of APAP, which was evident for inducing ER stress-related liver damage [12]. In context, worse liver function and survival rate and extensive hepatocyte death were manifested in IL-24-deficient group (Expanded View Fig. 3C-E). Given the possibility that the extracellular IL-24 might be implicated in liver injury, we treated WT mice with recombinant IL-24 (rIL-24) one hour before administration with CCL_4_. Nonetheless, the levels of transaminases, percentages of hepatocyte death and expression of P-PERK and CHOP showed no statistical differences between vehicle and cytokine-treated mice (Expanded View Fig. 4A-C).

**Fig. 2:**
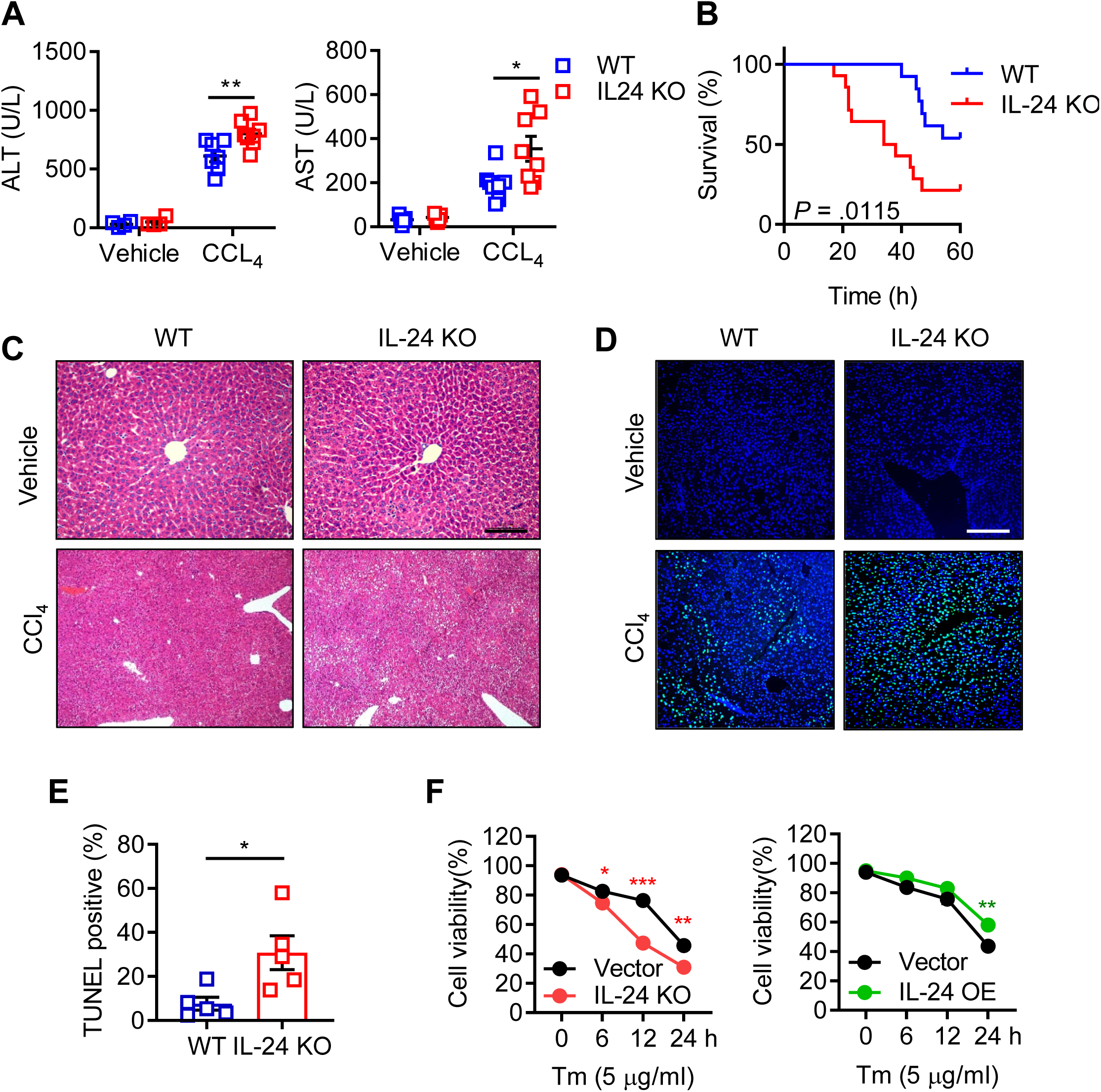
The protective role of hepatocyte IL-24 in ER stress-induced cell death. Sex- and age-matched WT and IL-24-null mice were intraperitoneally injected with vehicle or CCL_4_. (A) Mouse liver function was assessed by measuring serum ALT (left) and AST (right) levels. *n* = 5-8. (B) Mouse survival rate after CCL_4_-treatment was determined via Log-rank (Mantel-Cox) analysis. *n* = 11-13. (C, D) H&E (C) and TUNEL (D) staining of the liver tissues from vehicle or CCL_4_-treated mice. *n* = 5-8 mice. Scale bar, 100 µm. (E) Hepatocyte apoptosis after CCL_4_-treatment was assessed by counting TUNEL positive cells. *n* = 5-8 mice. (F) AML12 cells were transfected with lentiviral vectors expressing IL-24-targeted sgRNA (left, referred to as IL-24 KO) or IL-24 cDNA (right, referred to as IL-24 OE). An empty vector was transfected as a negative control, respectively. Cell viability of indicated AML12 cells after Tm stimulation was assessed by CCK8 assay. *n* = 3 biological replicates. Data are presented as means ± SEM. \**P* < 0.05, \*\**P* < 0.01, \*\*\**P* < 0.001. *P*-values were determined by two tailed *t*-test.

To obtain a closer insight into the intrinsic IL-24 function, we genetically depleted IL-24 in AML12 cells by using clustered regularly interspersed short palindromic repeats (CRISPR) / CRISPR associated protein 9 (Cas9) strategy and treated with Tm to mimic the pathological process *in vivo*. Cell Counting Kit-8 (CCK8) assays indicated that loss of intrinsic IL-24 impaired cell viability. Reciprocally, overexpression of IL-24 in AML12 benefited cell survival upon ER stress (Fig. 2F). Furthermore, annexin-V and propidium iodide staining showed that IL-24 attenuated late phase of apoptosis (Expanded View Fig. 5A&B). Together, these results suggested that hepatocyte IL-24 plays a fundamental role in protecting ER stress-mediated cell death.

### Hepatocyte IL-24 deficiency activates PERK-eIF2α-CHOP pathway

To understand the mechanism behind hepatocyte stress, we assessed CHOP expression in both two acute liver injury models and Tm-exposed AML12 cells. Remarkably, loss of IL-24 in the mouse liver or hepatocytes unleashed CHOP expression, while introduction of IL-24 into AML12 efficiently diminished CHOP level (Fig. 3A&B and Expanded View Fig. 6A&B&C). In line with these findings, IL-24 depletion upregulated the expression of pro-apoptotic factors such as Bim and TRIB3, and yet downregulated anti-apoptotic molecule Bcl2 (Expanded View Fig. 6D). To analyze how IL-24 was linked to ER stress, we further examined the UPR branches in the upstream of CHOP. Unexpectedly, either IRE1α phosphorylation or ATF6 expression appeared no difference between WT and IL-24 KO mice (Expanded View Fig. 6E). Noticeably, phosphorylation of PERK was selectively upregulated in the IL-24-deficient mouse liver or AML12 cells, and conversely downregulated upon IL-24 overexpression (Fig. 3A&B and Expanded View Fig. 6A). Accordingly, we evaluated the downstream molecules of PERK and found IL-24 deficiency robustly reinforced phosphorylation of eIF2α and expression of ATF4 and GADD34 in the stressed liver or AML12 cells, while overexpression of IL-24 in AML12 cells inhibited these molecules (Fig. 3A-E and Expanded View Fig. 6A-C). Immunohistochemical staining further confirmed higher levels of P-eIF2α and CHOP in IL-24-deficient liver (Fig. 3C). In aligned with these results, primary mouse hepatocytes isolated from IL-24-null mice manifested excessive activation of PERK-CHOP signal upon Tm-induced unresolved ER stress (Fig. 3F). Similarly, we also confirmed the activation of PERK-CHOP pathway in IL-24-deficient cells under *in vitro* CCL_4_ treatment (Expanded View Fig. 6F).

**Fig. 3:**
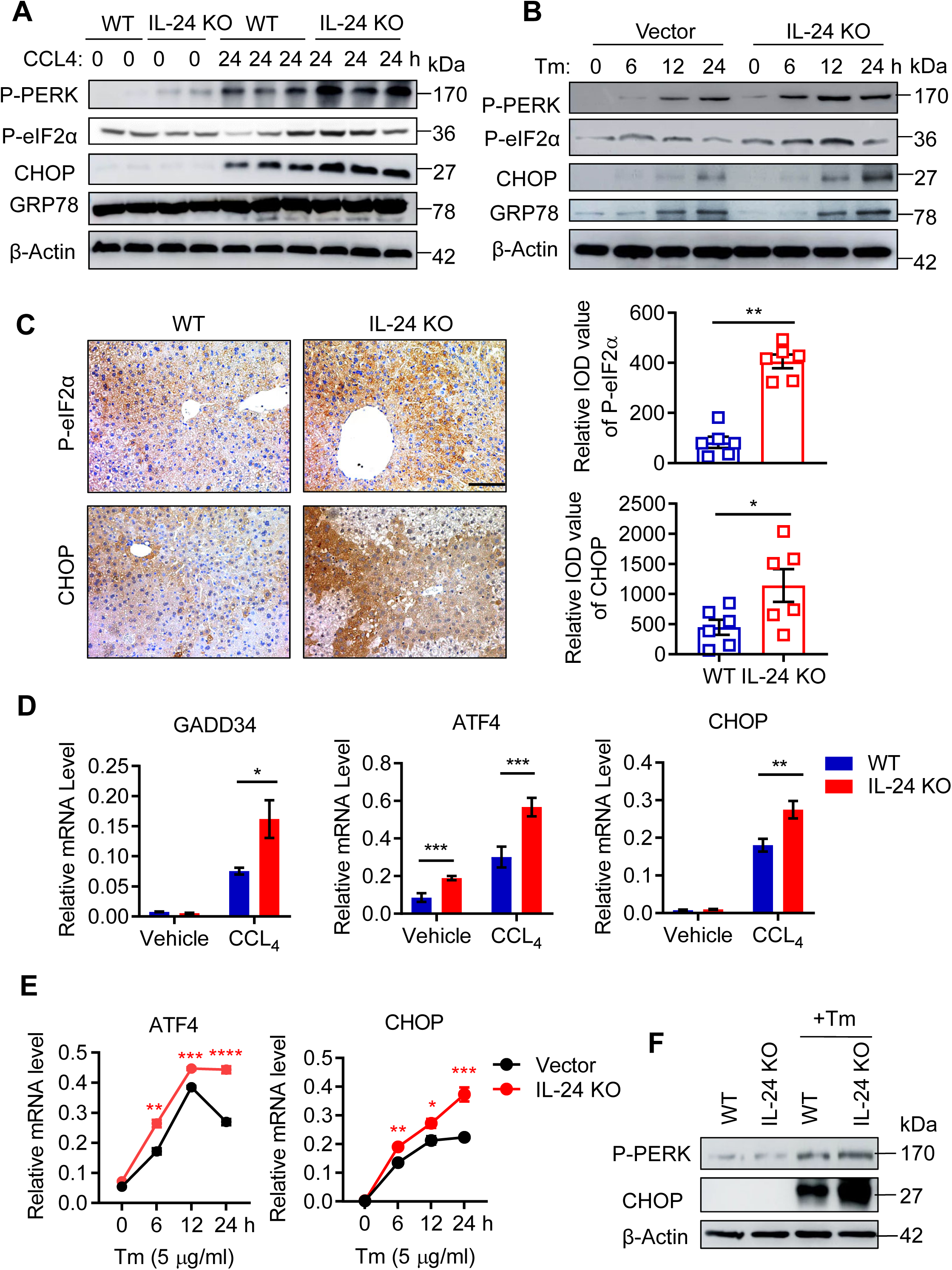
IL-24 deficiency facilitates PERK-eIF2α-CHOP UPR in hepatocyte. (A, B) P-PERK, P-eIF2α, CHOP and GRP78 protein levels in CCL_4_-treated WT and IL-24 KO mice (A) and Tm-stimulated control and IL-24 KO AML12 cells (B), as evaluated by western blot at indicated time points. *n* = 3 independent experiments. (C) Immunohistological staining of P-eIF2α and CHOP in the liver tissues from CCL_4_-treated WT and IL-24 KO mice. *n* = 5-8 mice. Scale bar, 100 µm. Results were represented in median integrated optical density (IOD) value. (D) GADD34, ATF4 and CHOP mRNA levels in the liver tissues from vehicle or CCL_4_-treated mice. *n* = 4-7 mice. (E) ATF4 and CHOP mRNA levels in indicated AML12 cells after Tm stimulation as assessed by qRT-PCR at indicated time points. (F) P-PERK and CHOP protein levels in Tm-stimulated primary hepatocytes from WT and IL-24 KO mouse, as evaluated by western blot. *n* = 3 independent experiments. Data are presented as means ± SEM. \**P* < 0.05, \*\**P* < 0.01, \*\*\**P* < 0.001, \*\*\*\**P* < 0.0001. *P*-values were determined by two tailed *t*-test.

### Hepatocyte IL-24 selectively limits CHOP-mediated death signal

To explore whether CHOP is indispensable for IL-24 deficiency-related hepatocyte damage, we utilized siRNA targeting CHOP (siCHOP) to comprehend its executive role in ER-stressed hepatocytes. Administration of siCHOP to AML12 cells offset the marginal cell death caused by IL-24 depletion (Fig. 4A). In addition, we crossed IL-24-null mice with a CHOP-null strain to generate a double knockout (DKO) strain. In contrast to IL-24 KO counterparts, both CHOP KO and DKO mice rejected to CCL_4_-induced liver injury (Fig. 4B). As evident in histological staining, disruption of CHOP eliminated hepatocyte death in IL-24 KO mice (Fig. 4C). Therefore, these results suggest that IL-24 deficiency promotes hepatocyte death dependently on CHOP in the context of unresolved ER stress.

**Fig. 4:**
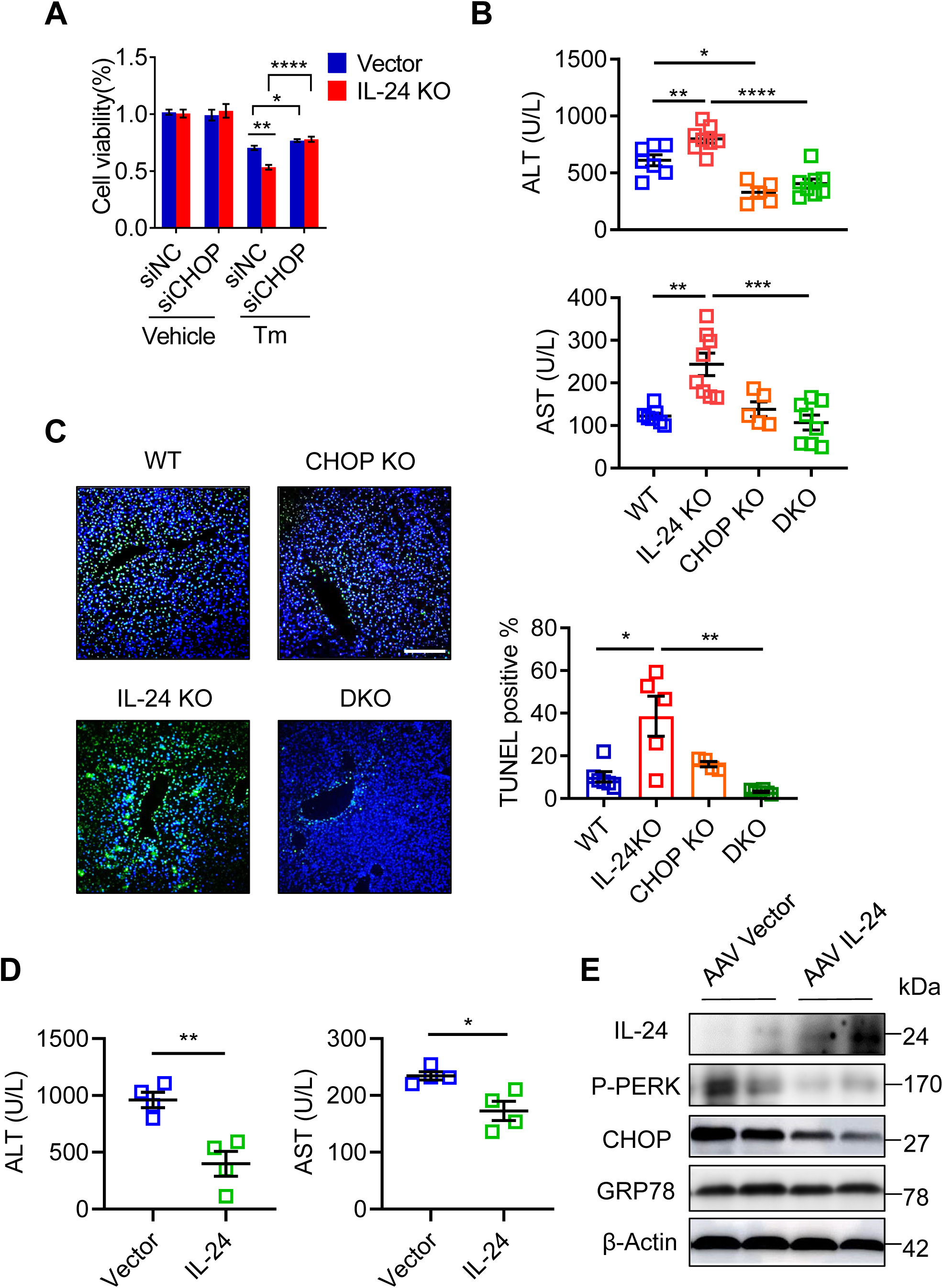
Hepatocyte IL-24 protects CHOP-executed cell death. (A) AML12 cells were treated with siCHOP or negative control (NC). Cell viability of indicated AML12 cells after Tm stimulation was assessed by CCK8 assay. *n* = 3 biological replicates. (B) Serum ALT (upper) and AST (lower) levels in WT, IL-24 KO, CHOP KO and IL-24/CHOP double KO (DKO) mice treated with CCL_4_ for 24 h. *n* = 5-8 mice. (C) TUNEL staining (left) of the liver tissues from CCL_4_-treated mice and quantification of TUNEL positive cells (right). Scale bar, 100 µm. (D) IL-24 KO mice were intravenously injected with AAV particles expressing an empty vector or mouse IL-24 8 weeks prior to CCL_4_ administration. Liver injury was assessed by serum ALT and AST levels. *n* = 4 mice. (E) IL-24, P-PERK, P-eIF2α, CHOP and GRP78 protein levels in the liver tissues from CCL_4_-exposed IL-24 KO mice with or without IL-24 re-expression. *n* = 3 independent experiments. Data are presented as means ± SEM. \**P* < 0.05, \*\**P* < 0.01, \*\*\**P* < 0.001, \*\*\*\**P* < 0.0001. *P*-values were determined by two tailed *t*-test.

To better understand the intrahepatic function of IL-24, we treated the IL-24 null mice with IL-24-expressing adeno-associated viral (AAV) particles 8 weeks prior to CCL_4_ administration. As shown in Fig. 4D&E, re-expression of IL-24 in the liver markedly reduced the levels of serum transferases and P-PERK, P-eIF2α and CHOP.

### Hepatocyte IL-24 attenuates liver damage by restricting the PERK-eIF2α branch

To ascertain the importance of PERK-eIF2α UPR branch, we built on an observation made with the inhibitor of the integrated stress response (ISRIB), which specifically blocks PERK-eIF2α signaling but unaffects ATF6 or inositol-requiring enzyme 1 (IRE1α) pathway[24]. Strikingly, pretreating AML12 with ISRIB compensated the viability loss for the lack of IL-24 but did not change the viability of WT cells (Fig. 5A). Next, we treated IL-24-null mice with vehicle or ISRIB one hour prior to CCL_4_ administration. As indicated by serum aminotransferases, ISRIB reversed the deterioration of liver damage in IL-24-null mice but showed no profound impact on WT mice (Fig. 5B). H&E and TUNEL staining visualized an amelioration in hepatocyte death intensified by IL-24 depletion (Fig. 5C), which could be explained by the reduction of CHOP expression after ISRIB treatment (Fig. 5D).

**Fig. 5:**
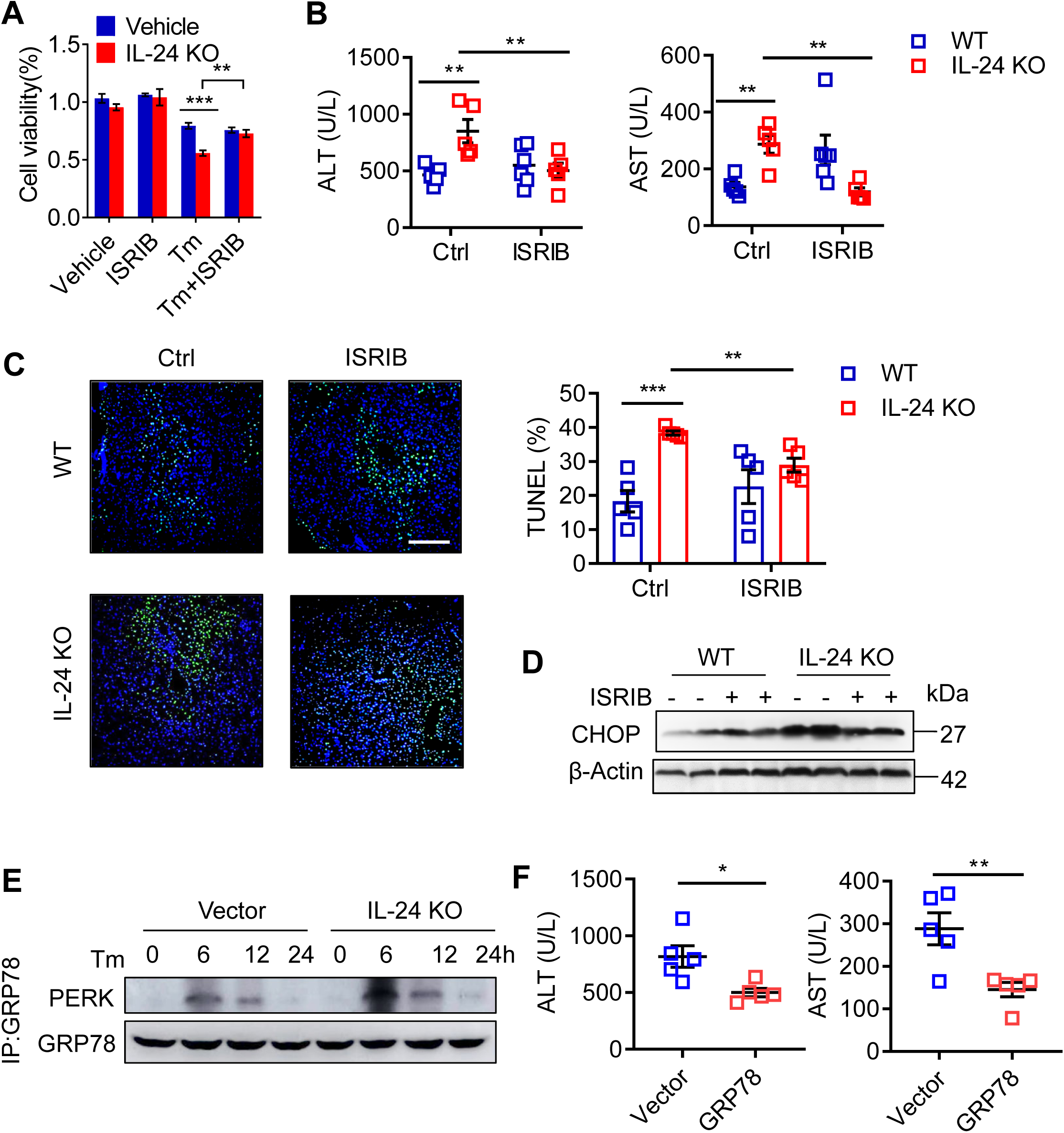
Perturbation of PERK-eIF2α signaling compensates ER homeostasis upon IL-24 loss. (A) Indicated AML12 cells were pre-treated with 200 nM ISRIB or DMSO (Vehicle) 24 h prior to Tm exposure. Cell viability of indicated AML12 cells after Tm stimulation was assessed by CCK8 assay. *n* = 3 biological replicates. (B, C) WT and IL-24 KO mice were intraperitoneally injected with vehicle or ISRIB (0.25 mg/kg) 60 min prior to CCL_4_ administration. Liver injury was assessed by serum ALT and AST levels (B) and TUNEL positive cell ratios (C). *n* = 5 mice. Scale bar, 100 µm. (D) Immunoblotting of CHOP in the liver tissues from CCL_4_-exposed mice with or without ISRIB treatment. *n* = 3 independent experiments. (E) Immunoblotting of PERK in the precipitates obtained by immunoprecipitation of endogenous GRP78 in indicated AML12 cells. *n* = 3 independent experiments. (F) IL-24 KO mice were intravenously injected with AAV particles expressing an empty vector or mouse GRP78 8 weeks prior to CCL_4_ administration. Liver injury was assessed by serum ALT and AST levels. *n* = 5 mice. Data are presented as means ± SEM. \**P* < 0.05, \*\**P* < 0.01, \*\*\**P* < 0.001. *P*-values were determined by two tailed *t*-test.

It is known that the ER protein chaperon GRP78 binds to the cytoplasmic and ER luminal domains of PERK to prevent its activation[25]. Accordingly, we asked whether hepatocyte IL-24 affected the interaction between GRP78 and PERK. By performing immunoprecipitation, we pulled down GRP78 in AML12 cells and detected a significant binding of PERK after 6 h of Tm stimulation. In concert with the UPR degrees, its association with PERK was significantly enhanced in IL-24-deficient cells and was weakened in IL-24-overexpressed cells (Fig. 5E and Expanded View Fig. 7A). To obtain a functional relevance *in vivo*, we replenished chaperon expression in the mouse liver by intravenous injection of GRP78-expressing AAV. As shown in Fig. 5F and Expanded View Fig. 7B&C, overexpression of GRP78 strongly mitigated liver damage and PERK-CHOP activation in the CCL_4_-treated IL-24-null mice.

### Hepatocyte IL-24 predicts prognosis for patients with acute liver injury

To examine the biological significance of IL-24 in clinical situations, we collected liver tissue and serum samples from 9 heathy donors, 9 patients with liver cirrhosis and 22 patients with acute liver failure (ALF). Serum IL-24 levels in both two groups of liver injury patients were as low as that in heathy donors (data not shown), which was aligned with what we found in CCL_4_-induced liver injury mouse models. Nonetheless, immunohistochemistry, immunoblots and qRT-PCR indicated that IL-24 expression was reduced in patients with liver cirrhosis and even lower in those with ALF (Fig. 6A-C&F), whereas CHOP expression was concomitantly escalated in cirrhosis and ALF patients (Fig. 6A&F), suggestive of a strong correlation between IL-24 expression and hepatocyte stress. In a further analysis of ALF patients, we obtained the individual liver function test before liver transplant and found IL-24 expression in hepatocytes was negatively related to serum ALT level as well as liver CHOP expression (Fig. 6D&E). Taken together, hepatocyte IL-24 may function as a prognosis predictor for patients with acute liver injury.

**Fig. 6:**
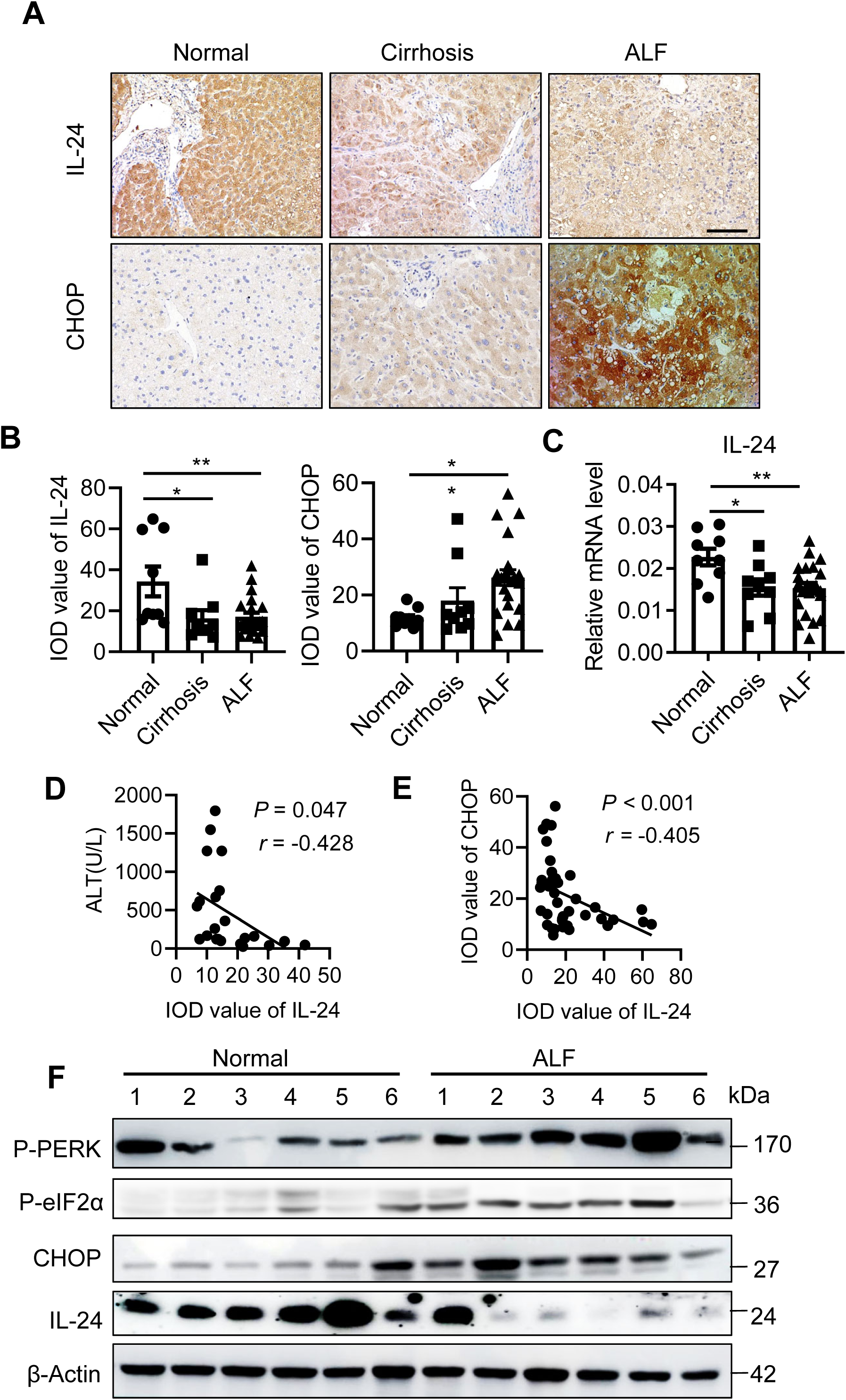
Hepatocyte IL-24 inversely correlates with liver function and CHOP expression in patients. (A) Immunohistological staining of IL-24 and CHOP in the liver tissues from healthy donors (*n* = 9), cirrhosis (*n* = 9) and acute ALF (*n* = 22) patients. Scale bar, 100 µm. (B) Quantification of IL-24 protein levels in (A), as represented in median integrated optical density (IOD) value. (C) IL-24 mRNA levels in the human liver tissues indicated in (A), as assessed by qRT-PCR. Results are normalized to β-actin. *n* = 9-22 patients. (D, E) Pearson correlation analysis between liver IL-24 protein level and serum ALT level (D) or liver CHOP protein level (E). *n* = 22. (F) Immunoblotting of IL-24, P-eIF2α and CHOP in the liver tissues from healthy donors and acute ALF patients. *n* = 9 patients. Data are presented as means ± SEM. \**P* < 0.05, \*\**P* < 0.01. *P*-values were determined by two tailed *t*-test.

**Fig. 7:**
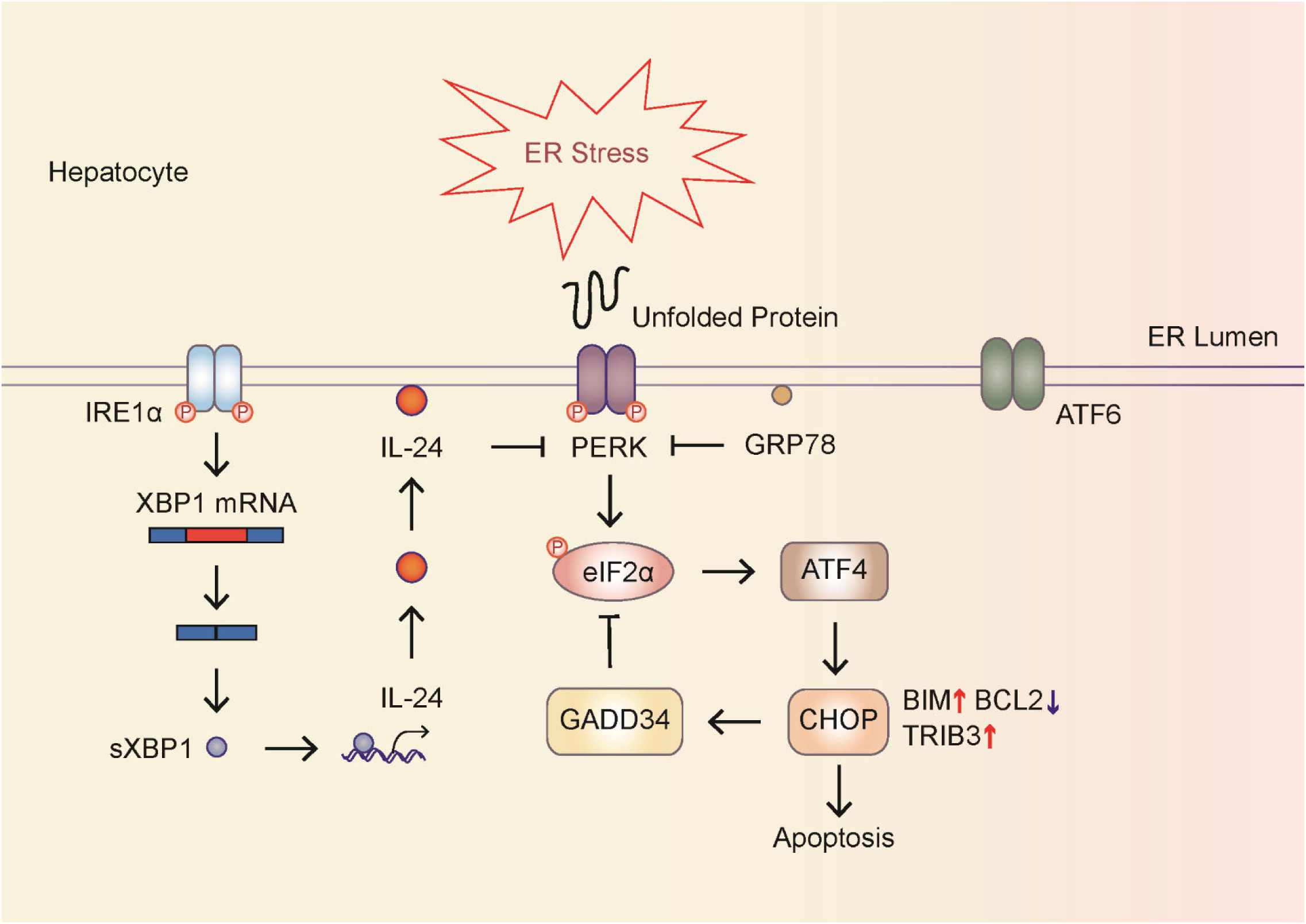
Schematic model showing the interaction between cytoplasmic IL-24 and UPR modulators within hepatocytes. Hepatocyte ER stress engages sXBP1 for upregulating IL-24 transcription, which in turn improves ER homeostasis and represses CHOP-mediated cell death by harnessing PERK-eIF2α branch reaction.

## Discussion

UPR is executed through three ER transmembrane stress sensors: IRE1α, PERK and activating transcription factor 6 (ATF6)[5]. Activated IRE1α splices XBP1 mRNA, which encodes transcription factor to increase the protein-folding capacity and degrade misfolded proteins. While IRE1α engages STAT3 pathway to promote liver regeneration upon liver injury [26], XBP1 switches pro-survival to pro-apoptotic signal cascades through multiple gene regulation [27, 28]. It has been reported that ATF6 exerts a pro-inflammatory effect on ischemia-reperfusion liver injury [29]. PERK antagonizes UPR by reducing the flux of protein translocating and phosphorylating eIF2α, a pervasive translation initiation factor, which inhibits ribosome assembly and translation. However, eIF2α selectively upregulates the transcription factor ATF4 and its downstream target CHOP. Sustained ER stress engages CHOP to enhance UPR and inflammation signaling and lead to apoptosis [22, 30]. The mouse models in our study manifested activation of three UPR sensors and CHOP. The latter was also found to be upregulated in human cirrhosis and ALF patients. Given the recovery of liver function in most cases, we hypothesized an unknown machinery restoring hepatocyte ER homeostasis.

IL-24 has been reported to cause ER stress-mediated apoptosis through a secretion-independent manner [19]. Unexpectedly, we detected abundant IL-24 expression in the mouse or human liver under normal condition. In the CCL_4_ model, hepatocyte IL-24 increases instantaneously and then returns to baseline as the liver function recovers. Nonetheless, serum IL-24 was undetectable in CCL_4_-treated mice and human cirrhosis and ALF patients, excluding its engagement as a “hepatokine” in UPR condition. Importantly, targeting ER stress-related transcription factors (ATF4, ATF6, XBP1 and CHOP) significantly reduced the mRNA level of hepatocyte IL-24. However, only silencing of XBP1 deprived IL-24 expression under either normal or deleterious circumstance. This reflects a physiological function of hepatic XBP1 and also raises a possibility that intrinsic IL-24 may regulate stress signals.

Since adenovirus-mediated overexpression causes protein synthesis overload and induces potential UPR, we employed gene knockout mice to improve our understanding of cytosolic IL-24 in ER stress and liver damage. Remarkably, we found IL-24-null mice were more sensitive to CCL_4_-induced liver injury than WT counterparts. While either ATF6 expression or IRE1α phosphorylation was unaffected, P-PERK, P-eIF2α, CHOP and GADD34 exhibited excessive expression in the IL-24-deficient mouse liver. In addition, our results showed elevated levels of Bim and TRIB3 and a reduced level of Bcl2 in IL-24-null mice, which might be a consequence of CHOP activation [3, 11]. Effects of hepatocyte IL-24 on PERK-eIF2α-CHOP branch and cell death were corroborated by depleting or introducing IL-24 in AML12 cells. Furthermore, knocking out CHOP in IL-24-null mice or knocking down CHOP in IL-24-null AML12 protected ER stress-associated hepatocyte damage. It has been reported that eIF2α phosphorylation acts as a central event sensitizing stressed cells to death [12, 27, 29]. Accordingly, we found that the extensive liver damage in IL-24-null mice could be alleviated by administration of ISRIB, which selectively reverses the effects of eIF2α phosphorylation [24]. Clinically, we observed a concomitant downregulation of IL-24 and upregulation of CHOP in the cirrhosis and ALF tissues as compared with the healthy liver. Taken together, these findings demonstrated the protective role of IL-24 in resolving hepatocyte ER stress is implemented by perturbation of PERK-eIF2α-CHOP pathway.

As one of the most important ER chaperons, GRP78 is essential in conjunction with misfolded proteins and is important for maintaining ER homeostasis [8, 31]. Given the reinforced ER stress in IL-24-null hepatocytes, GRP78 conserved its affinity to combine and sequester overreacted PERK. Despite this, redundant PERK phosphorylated itself and triggered the downstream signaling. Previous study indicated that *in vivo* overexpression of GRP78 using an adenovirus vector could attenuate ER stress-associated liver steatosis [32]. In this study, introduction of GRP78 in the IL-24-deficient mouse liver by AAV infection attenuated PERK-facilitated hepatocyte stress. This may provide a potential therapeutic opportunity for UPR-related human liver diseases, especially those with low hepatocyte IL-24 expression.

To our knowledge, a variety of cytokines, including those expressed or secreted by hepatocytes, evoke inflammatory responses and promote cell death in liver diseases. In a diet-induced steatohepatitis mouse model, hepatocyte IL-1α was found to be upregulated in response to ER stress, which in turn enhanced CHOP expression; IL-1α released from necrotic hepatocytes accelerates steatohepatitis via induction of inflammatory cytokines [33, 34]. In lipopolysaccharide (LPS)-induced liver injury, hepatocyte-derived IL-7 augmented CD8+ T cell cytotoxic activity and promoted the development of autoimmune diseases [35]. In the present study, we showed that intracellular IL-24 uniquely benefited hepatocyte ER homeostasis, exerting an anti-inflammatory effect. However, it has not been defined whether hepatocyte can secrete IL-24 under pathological conditions and how it affects hepatocytes or immune cell populations in an autocrine or paracrine fashion.

Conclusively, we uncovered that cytosolic IL-24 is critical for protecting ER stressed-hepatocytes from death, which may be a good diagnostic and therapeutic target for clinical liver diseases.

## Materials and Methods

### Patients

Cirrhosis and liver failure tissues were collected from liver transplant recipients treated in Department of Liver Surgery, Renji Hospital, School of Medicine, Shanghai Jiaotong University. Normal liver tissues were collected from the healthy transplant donors through liver biopsy. All samples were collected with informed consent, and the experiments were approved by the ethical review committee of the World Health Organization Collaborating Center for Research in Human Production (authorized by the Shanghai Municipal Government).

### Mice

All mice used in this study were in C57BL/6J background. IL-24 knockout (KO) mice were generated in ShanghaiTech University. The targeted Embryonic stem (ES) cells were ordered from The Knockout Mouse Project (KOMP) Repository, in which the insertion of Velocigene cassette ZEN-Ub1 created a deletion of size 3306 bp between positions 132779010-132782315 of Chromosome 1 (Genome Build37). These ES cells were injected into albino C57BL/6J blastocysts and the following inbred strain was generated by backcrossing breeding. The following PCR primers were used to identify WT (571 bp) and KO (338 bp) alleles: 5’-GTACCCACTCCAATGCATACATT −3’, 5’-GCTCATCCAGGATGAAGCTACAC −3’, and 5’-GAAACCAGGCAAATCTCCACTCC −3’. The CHOP KO mice and Alb^Cre^ transgenic mice were purchased from Jackson Laboratories. We crossed the CHOP KO and IL-24 KO strains to produce the double knockout (DKO) strain. The Xbp1^F/F^ strain, a gift from Dana Farber Cancer Institute, USA, was previously described [36]. Mice devoid of XBP1 selectively in hepatocytes were generated by breeding the Xbp1^F/F^ mice with Alb^Cre^ strain. For acute liver injury model, CCL_4_ dissolved in olive oil was injected intraperitoneally into 8-week-old female mice at a dose of 2 ml/kg [26]. Besides, mice were fasted overnight and administered 500 mg/kg APAP (Sigma) by oral gavage [12]. In some settings, mice were intraperitoneally injected with 0.25 mg/kg ISRIB (Selleck) or recombinant IL-24 (R&D) at a dose of 5 μg per mouse one hour prior to CCL_4_ administration. All mice were maintained under specific pathogen-free (SPF) conditions, on a 12 h light-dark cycle. All mouse experiments were approved by the Shanghai Administrative Committee for Laboratory Animals.

### Isolation of primary mouse hepatocytes and cell cultures

The mouse liver was perfused with an EGTA-buffer (37°C) at a constant flow of 5 ml/min for 8 minutes via the hepatic portal vein. Next, a secondary perfusion with a solution of collagenase I (Sigma-Aldrich) for 10 minutes (2 ml/min) is required for a completely digestion. The liver was disrupted gently to release hepatocytes into a suspension buffer. Subsequently, the liver capsule was filtered through a 70-µm cell strainer and the primary hepatocytes were collected after three cycles of centrifugations at 400 rpm for 5 min at 4°C. A suspension of 1×10^6^ cells/mL was successively seeded in culture plates. Murine hepatocyte cell line AML12 was from The First Affiliated Hospital of Nanjing Medical University. Primary hepatocytes and AML12 cells were cultured in William’s E Medium (Gibco) supplemented with 1X insulin-transferrin-selenium supplement (Gibco), 1X sodium pyruvate (Gibco), 40 ng/ml dexamethasone (Sigma-Aldrich) and 10% FBS (Gibco), incubated at 37 °C and 5% CO_2_, and were tested for mycoplasma contamination once every three months. In some settings, cells were treated with 5 μg/ml Tm (Sangon Biotech) or 200 nM ISRIB (Selleck).

### siRNA, sgRNA and gene transfection

The siRNAs were transiently transfected into AML12 cells by using Lipofectamine RNAiMAX (Invitrogen) following manufacturer’s instructions. A non-target siRNA was used as negative control. The siRNA sequences are listed as below: mouse XBP1, CCAAGCUGGAAGCCAUUAATT; mouse CHOP, CCAGAUUCCAGUCAGAGUUTT; mouse ATF4, CUCCCAGAAAGUUUAAUAATT; mouse ATF6, GCAGUCGAUUAUCAGCAUATT. Stable knockout of IL24 in AML12 cells was generated by lentiviral-based delivery of sgRNA/cas9 components. Briefly, sgRNA targeting the exonic region of murine *Il24* gene (5′-GAAGGATTAGGCTCAGGCAG-3′) were subcloned into the lentiviral vector GV393 (U6-sgRNA-EF1a-Cas9-FLAG-P2A-EGFP) (Genechem, China), while a non-target sgRNA (5′-CGCTTCCGCGGCCCGTTCAA-3′) was used as a negative control. IL-24-overexpression construct was generated by subcloning PCR-amplified full-length cDNA (NM_053095) into a GV358 (Ubi-MCS-Flag-SV40-EGFP-IRES-puromycin) lentiviral vector (Shanghai Genechem). An empty vector was used as a negative control. Viral particles were packaged in 293T cell and used to infect AML12 cells in the presence of 8 µg/ml polybrene followed by puromycin selection.

### Recombinant AAV construction and In vivo transduction

GRP78-overexpression construct was generated by subcloning PCR-amplified full-length *Il24* (NM_053095) or *Hspa5* (NM_022310) cDNA into a GV461 (CMV-betaGlobin-MCS-SV40 PolyA) AAV vector (Shanghai Genechem). An empty vector was used as a negative control. IL24 KO mice of 5 weeks were intravenously injected with 2 × 10^11^ vector-genome (vg) AAV 8 weeks prior to CCL_4_ administration.

### Immunohistochemistry and Immunofluorescence

Immunohistochemistry for target molecules was performed on serial sections from human or mouse liver tissues. Sections were deparaffinized, subjected to antigen retrieval, and incubated with primary antibodies against IL-24 (Abcam, ab115207), P-eIF2a (Huabio, ET1603-14) and CHOP (Huabio, ET1703-05). All responses were followed by staining with the corresponding HRP-conjugated secondary antibody (Jackson Immuno Research Laboratories). The stained slides were assessed with integrated optical density (IOD) using ImageJ software. The apoptotic cells were defined by using In Situ Cell Death Detection Kit (Roche) following manufacturer’s protocol and quantified by calculating positively stained cells in at least five randomly chosen HPFs of each slide.

### Immunoprecipitation

Cell samples were collected and lysed in IP lysis buffer (Thermo Fisher) containing protease inhibitor cocktail (Merck Millipore) for 30 minutes. After an insoluble product-clear step by full speed centrifuge, the supernatant was harvest and incubated with anti-GRP78 (Abcam, ab21685) antibody and protein A beads (Thermo Fisher) at 4 °C overnight. The beads were collected and washed extensively, and the immuno-complex was eluted with western blot loading buffer.

### Western blot

Cell or tissue lysates were separated on 6-8% polyacrylamide-SDS gels and transferred to a nitrocellulose membrane using transfer buffer (25 mM Tris, 192 mM glycine and 10% methanol). The blots were blocked with 5% non-fat milk in PBS containing 0.05% Tween-20 for 1 h and then probed overnight at 4°C in PBST with primary antibodies. Next, the blots were incubated with a secondary antibody conjugated to horseradish peroxidase (HRP) (1:5,000, Jackson Immuno Research Laboratories) and detected with a ChemiDoc XRS system (Bio-Rad). Primary antibodies used in western blot are listed below: anti-Flag (14793S), anti-P-PERK (3179S), anti-ATF4 (11815S) and anti-CHOP (2895S) were from Cell Signal Technology, anti-human IL-24 (ab115207), anti-P-eIF2a (ab32157) and anti-GRP78 (ab21685) were from Abcam, anti-mouse IL-24 (MAB2786) was from R&D Systems, anti-PERK (sc-377400) was from Santa Cruz.

### Quantitative PCR (qPCR)

Total RNA samples used for RT-qPCR were isolated by using an RNeasy kit (BioTeke) with an additional on-column DNase-I digestion step. Total RNA or purified mRNA was reverse transcribed with PrimeScript™ RT Master Mix (Takara) using Oligo dT primers to obtain complementary DNA. qPCR was carried out by using SYBR Premix Ex Taq II (Takara). β-actin was used as an internal control. The primers used in this study are: mouse GRP78(F:5’-TCATCGGACGCACTTGGAA −3’;R:5’-CAACCACCTTGAATGGCAAGA −3’); mouse CHOP(F:5’-CTGGAAGCCTGGTATGAGGAT-3’;R:5’-CAGGGTCAAGAGTAGTGAAGGT −3’) ; mouse ATF4(F:5’ - CTCTTGACCACGTTGGATGAC - 3’; R: 5’ - CAACTTCACTGCCTAGCTCTAAA −3’); mouse IL-24 (F:5’- GAGCCTGCCCAACTTTTTGTG −3’;R:5’- TGTAGTCCCCAACTCATCTGTG −3’); mouse sXBP1 (F: 5’- CTGAGTCCGAATCAGGTGCAG −3’; R: 5’- GTCCATGGGAAGATGTTCTGG - 3’); mouse ATF6 (F: 5’- TCGCCTTTTAGTCCGGTTCTT −3’; R: 5’- GGCTCCATAGGTCTGACTCC - 3’); mouse GADD34 (F: 5’- GAGGGACGCCCACAACTTC −3’; R: 5’- TTACCAGAGACAGGGGTAGGT - 3’); mouse IL6 (F: 5’- TAGTCCTTCCTACCCCAATTTCC −3’; R: 5’- TTGGTCCTTAGCCACTCCTTC - 3’); mouse IL1A (F: 5’- CGAAGACTACAGTTCTGCCATT −3’; R: 5’- GACGTTTCAGAGGTTCTCAGAG - 3’); mouse Bim (F: 5’- GACAGAACCGCAAGGTAATCC −3’; R: 5’- ACTTGTCACAACTCATGGGTG - 3’); mouse TRIB3 (F: 5’- GCAAAGCGGCTGATGTCTG −3’; R: 5’-AGAGTCGTGGAATGGGTATCTG - 3’); mouse Bcl2 (F: 5’- ATGCCTTTGTGGAACTATATGGC −3’; R: 5’- GGTATGCACCCAGAGTGATGC - 3’); mouse TNFA (F: 5’- CTCTTCTGTCTACTGAACTTC −3’; R: 5’- CTCCTGGTATGAGATAGCAA - 3’); mouse β-actin (F:5’- ACCCACACTGTGCCCATCTAC −3’;R:5’- AGCCAAGTCCAGACGCAGG −3’); human IL-24(F:5’- CACACAGGCGGTTTCTGCTAT-3’; R:5’- TCCAACTGTTTGAATGCTCTCC −3’); human β-actin (F: 5’- GGGAAATCGTGCGTGACATTAAG −3’; R: 5’- TGTGTTGGCGTACAGGTCTTTG - 3’).

### Dual luciferase assay

A DNA fragment of *Il24* (−1036 ∼ −598 bp in the upstream of transcription start site) was subcloned into a luciferase reporter vector pGL4 (Promega). AML12 cells were cultured in 24- well plates for 24 hours, then transfected with siRNAs. The cells were co-transfected with luciferase reporter plasmid and renilla luciferase plasmid (an internal control) at a ratio of 10:1. Twenty-four hours later, cells were treated with Tm (5 μg/ml). Cells from each independent well were harvested and detected by Dual-Luciferase Reporter Assay System (Promega) at indicated time. Get the relative light unit (RLU) by normalizing to renilla luminescence activities.

### Apoptosis analysis

AML12 cells were stimulated by Tm, and 1∼5×10^5^ cells were collected by centrifugation. The cells were resuspended with the 1× Binding Buffer and incubated with 5μL FITC-conjugated annexin V Annexin FITC (BD Biosciences, USA) and 5μL PI (BD Biosciences, USA) to each tube according to the experimental protocol. Then samples were analyzed by fluorescence-activated cell sorter (FACS).

### Statistical analysis

At least three biological replicates were used in each experiment unless otherwise stated. Data were analyzed with GrapPad Prism 7 and were presented as the mean ± standard error of the mean (SEM). Two-tailed Student’s t-tests were performed to assess the statistical significance of differences between groups. Pearson correlation coefficients (*r*) were calculated to assess correlation and statistical significance was assessed by a two-tailed t-test of *r* = 0.

### Expanded View

for this article is available online

## Acknowledgements

We thank all patients and donors for participating in this study. We thank Ms Jiaxin Li (Renji Hospital, Shanghai, China) for helping with clinical sample collection, Dr. Indrajit Das (QIMR Berghofer Medical Institute, QLD, Australia) for the advice on ER stress antagonism, and Dr. Laurie H. Glimcher (Dana-Farber Cancer Institute, MA, USA) for providing Xbp1^F/F^ mice as a kind gift. This study was supported by National Natural Science Foundation of China (81672801 to JH, 81670598 to QX and 81700498 to Haiyan Z), Chen Guang Project in Shanghai Municipal Education Commission and Shanghai Education Development Foundation (15CG13 to JH and 17CG20 to Haiyan Z) and National Key Research and Development Program of China (2016YFC0905901 to XH).

## Author contributions

J.H., X.H. and Q.X. conceived and designed the project, analysed and interpreted results, obtained funding and wrote the manuscript; J.W. and B.H. designed and performed experiments, analysed and interpreted results and wrote the manuscript; He Zhang, Zhenjun Zhao, Zhicong Zhao contributed to design and conduction of experiments and to analysis and interpretation of data; Haiyan Zhang, XM, Bin Shen and Beicheng Sun contributed to conception of research.

## Expanded View Figure Legends

**Expanded View Fig. 1 Determination of the levels of IL-24 and UPR markers during hepatocyte stress.** (A) Protein levels of IL-24 in different mouse tissues as assessed by western blot. *n* = 3 independent experiments. (B) Mouse liver function was evaluated by measuring serum ALT (left) and AST (right) levels 0-72 h post CCL_4_ injection. *n* = 4. (C, D) ATF4, ATF6 and sXBP1 mRNA levels in CCL_4_-treated mice (C) and Tm-stimulated AML12 cells (D), as evaluated by qRT-PCR at indicated time points. *n* = 3 biological replicates. Data are presented as means ± SEM. \**P* < 0.05, \*\**P* < 0.01, \*\*\**P* < 0.001, \*\*\*\**P* < 0.0001. *P*-values were determined by two tailed *t*-test.

**Expanded View Fig. 2 Hepatocyte XBP1 is essential for maintaining IL-24 transcription.** (A) Alignment of responsive elements for CHOP (boxed) XBP1/ATF6 (red) found in the promoter region of *Il24*. The base positions of the consensus are indicated 5′→3′. Positions are relative to transcription initiation site. (B) AML12 cells were transfected with indicated siRNAs or negative control (NC) 24 h prior to Tm treatment. IL-24 mRNA levels at indicated time points were assessed by qRT-PCR. (C) *Il24* promoter activity in AML12 cells expressing siRNAs with or without Tm treatment (24 h), as quantified using luciferase assay. *Renilla* luciferase activity was normalized to firefly activity and presented as relative luciferase activity. *n* = 3 biological replicates. (D) Cell viability of Ctrl (Xbp1^f/w^) and XBP1 KO (Xbp1^f/w^Alb^Cre^) mouse hepatocytes at indicated time points post Tm treatment was assessed by CCK8 assay. *n* = 3 independent experiments. (E, F) Protein (E) and mRNA (F) levels of IL-24 in Ctrl (Xbp1^f/w^) and XBP1 KO (Xbp1^f/w^Alb^Cre^) mouse hepatocytes at indicated time points post Tm treatment. *n* = 3 independent experiments (E) or biological replicates (F). Data are presented as means ± SEM. \**P* < 0.05, \*\**P* < 0.01, \*\*\**P* < 0.001, \*\*\*\**P* < 0.0001. *P*-values were determined by two tailed *t*-test.

**Expanded View Fig. 3 IL-24 deficiency exacerbates ER stress-related liver injury.** (A) PCNA staining of the liver tissues from CCL_4_-treated WT and KO mice. Hepatocyte proliferation after CCL_4_-treatment was assessed by counting PCNA positive cells. Scale bar, 100 µm. (B) IL6, TNFA and IL1A mRNA levels in the liver tissues from vehicle or CCL_4_-treated WT and IL-24 KO mice. *n* = 3 mice. (C-E) Sex- and age-matched WT and IL-24-null mice were orally treated with vehicle or APAP (500 mg/kg). (C) Mouse liver function was assessed by measuring serum ALT (left) and AST (right) levels. *n* = 5-8. (D) Mouse survival rate after APAP-treatment was determined via Log-rank (Mantel-Cox) analysis. *n* = 9-10. (E) H&E and TUNEL staining of the liver tissues from APAP-treated WT and IL-24-null mice. *n* = 5-8. Scale bar, 100 µm. Data are presented as means ± SEM. \**P* < 0.05, \*\**P* < 0.01, \*\*\**P* < 0.001. *P*-values were determined by two tailed *t*-test.

**Expanded View Fig. 4 Extracellular IL-24 does not affect liver function.** Recombinant IL-24 (rIL-24) (5 μg per mouse) was intraperitoneally treated one hour prior to CCL_4_ administration. (A) Mouse liver function was assessed by measuring serum ALT (left) and AST (right) levels. *n* = 6 mice. (B) H&E and TUNEL staining of the liver tissues from Vehicle or rIL-24-pre-treated mice. Scale bar, 100 μm. (C) P-PERK and CHOP protein levels in Vehicle or recombinant IL-24-pre-treated mice as evaluated by western blot at indicated time points. *n* = 3 independent experiments. Data are presented as means ± SEM. \*\**P* < 0.01, \*\*\**P* < 0.001. *P*-values were determined by two tailed *t*-test.

**Expanded View Fig. 5 Intrinsic IL-24 reduces Tm-induced hepatocyte apoptosis.** (A, B) Ratios of early and late phases of apoptosis in AML12 cells expressing different levels of IL-24 with or without Tm treatment, as evaluated by Annexin V-PI staining. Data are presented as means ± SEM. \**P* < 0.05, \*\*\**P* < 0.001, \*\*\*\**P* < 0.0001. *P*-values were determined by two tailed *t*-test.

**Expanded View Fig. 6 Hepatocyte IL-24 attenuates PERK-eIF2α-CHOP branch reaction.** (A) P-PERK, P-eIF2α, CHOP and GRP78 protein levels in Tm-stimulated control and IL-24 OE AML12 cells, as assessed by western blot. *n* = 3 independent experiments. (B) CHOP (upper) and ATF4 (lower) mRNA levels in Tm-stimulated control and IL-24 OE AML12 cells, as evaluated by by qRT-PCR at indicated time points. *n* = 3 biological replicates. (C) CHOP and GRP78 protein levels in APAP-treated WT and IL-24 KO mice, as evaluated by western blot at indicated time points. *n* = 3 independent experiments. (D) Bim, TRIB3 and Bcl2 mRNA levels in the liver tissues from vehicle or CCL_4_-treated WT and IL-24 KO mice. *n* = 3 independent experiments. (E) P-IRE1α and ATF6 protein levels in CCL_4_-treated WT and IL-24 KO mice, as evaluated by western blot at indicated time points. *n* = 3 independent experiments. (F) P-PERK and CHOP protein levels in CCL_4_-treated AML12 cells, as evaluated by western blot at indicated time points. *n* = 3 independent experiments. Data are presented as means ± SEM. \*\**P* < 0.01, \*\*\**P* < 0.001. *P*-values were determined by two tailed *t*-test.

**Expanded View Fig. 7 GRP78 compensates the anti-stress function of hepatocyte IL-24.** (A) Immunoblotting of PERK in the precipitates obtained by immunoprecipitation of endogenous GRP78 in indicated AML12 cells. *n* = 3 independent experiments. (B) IL-24 KO mice were intravenously injected with AAV particles expressing an empty vector or mouse GRP78 8 weeks prior to CCL_4_ administration. Liver injury was assessed by TUNEL positive cell ratios. *n* = 5 mice. Scale bar, 100 µm. (C) Immunoblotting of P-PERK and CHOP in the liver tissues from CCL_4_-exposed IL-24 KO mice with or without GRP78 overexpression. *n* = 3 independent experiments. Data are presented as means ± SEM. \*\**P* < 0.01. *P*-values were determined by two tailed *t*-test.

